# Convergent biology, divergent drivers: a cross-species comparison of human and canine invasive urothelial carcinoma

**DOI:** 10.64898/2026.08.05.742970

**Authors:** Haejin Cho, Jonathan P. Mochel, Megan P. Corbett, Lilian J. Oliveira, Karin Allenspach, Christopher Zdyrski, Aleksandra Pawlak, Burles A. Johnson, Eugene F Douglass

## Abstract

Traditional animal models are often inbred and genetically uniform. This makes them powerful for controlled experiments, but it limits how well they represent the patient-to-patient variation seen in real-world disease. Comparative oncology seeks to address this gap by studying naturally occurring cancers in outbred companion animals, especially dogs. Canine medicine offers two important advantages: first, prospective trials can often be completed faster than in humans and second, dogs are already part of the translational pipeline through pharmacokinetic and toxicology studies. Here, we assessed the transcriptional fidelity of human and canine invasive urothelial carcinoma in primary tumors and patient-derived organoids. We then used single-cell and spatial data to resolve the underlying cellular organization.

Despite strong species and platform differences, human and canine tumors preserved the same major luminal-basal structure and a similar tumor microenvironment. The two species reached this shared biology through different recurrent mutations. These included FGFR3 alterations in humans and BRAF alterations in dogs, which converged on overlapping pathways and a luminal phenotype. Human and canine organoids also underwent a similar shift in culture. Both became more proliferative and metabolic while losing inflammatory programs. Thus, organoids preserved important tumor biology while introducing predictable platform effects. Single-cell and spatial analyses showed that the luminal-basal axis reflects a gradient of cell states organized around the tumor-stroma boundary, rather than two discrete tumor types. This helps explain why bulk RNA-sequencing subtypes are reproducible but coarse. Together, these findings define where canine and human bladder cancer agree, where they differ, and how dogs can support parallel therapeutic and diagnostic development.

## INTRODUCTION

Animal models are fundamental to biomedical research because they enable experiments that are impractical or impossible in human patients.^1, 2^ Their value extends into clinical studies as well: having shorter lifespans, accelerated disease progression, and lower costs allow statistically powered studies of disease mechanisms and therapeutic response to be completed on timescales that would require years in human patients.^3, 4^ The challenge is translation, where only 5% of therapeutic interventions validated in animal models obtain regulatory approval in humans.^5^ For discoveries made in animals to inform human biology, researchers must distinguish features that are conserved across species from those that diverge.^6^ Currently, the identification of candidate therapeutics appropriate for clinical trials utilizing current pre-clinical research methods is poor, as only 3-5% of drugs that enter phase I oncology clinical trials,^7^ and 16-20% of drugs in phase II studies,^8^ are ever FDA approved for human cancer therapy. Thus, increasing fidelity and applicability of pre-clinical studies to human disease is critical to improve patient outcomes, while also preserving the scarce resource of patients appropriate for clinical trials only for drugs have a high chance of success.

However, conservation alone is not sufficient for translational fidelity, because human disease is defined by substantial patient-to-patient heterogeneity. Many traditional laboratory rodent models are highly inbred, and this genetic uniformity, removes much of the biological variation that underlies patient- heterogeneity underlying molecular diagnostics.^9, 10^ Model fidelity therefore depends on three related questions: what biology is *<u>conserved across species</u>* (C), what *<u>diverges across species</u>* (D), and whether clinically relevant *<u>patient-to-patient heterogeneity</u>* (H) is preserved.

One Health initiatives seek to address these limitations by leveraging naturally occurring diseases in veterinary species to complement traditional laboratory models.^11^ Programs such as the NCI Comparative Oncology Program recognize that companion animals develop spontaneous cancers within intact immune systems, shared environments, and outbred populations that more closely resemble human patients than laboratory rodents.^4^ Importantly, this rationale is not limited to cancer, as dogs also develop spontaneous cardiovascular, cardiorenal-metabolic, gastrointestinal, ophthalmic, neurologic, and endocrine diseases with conserved features relevant to human physiology and drug development.^11,12^ Consequently, veterinary cohorts capture inter-patient heterogeneity comparable to that observed in human populations, enabling evaluation of both average biological behavior and patient-level variation.^13, 14^ Canine studies are particularly attractive for two reasons: first, prospective clinical trials are feasible at a scale and speed difficult to achieve in humans;^15^ and second, dogs are already embedded in human translational research through their use in early pharmacokinetic and toxicology studies.^5^ *Moreover, many standards of care are shared across veterinary and human oncology, creating a bidirectional relationship in which advances in one species can inform the other*.^3, 4^

Bladder cancer (BC) is extensively characterized in both humans and dogs, lending itself well for cross-species comparison.^9, 13, 16–18^ In humans, muscle invasive bladder cancer (MIBC) is a leading cause of urologic cancer death.^19^ Large transcriptomic efforts have converged on a two-tier classification in which *luminal* and *basal* molecular subtypes capture the dominant axis of inter-patient heterogeneity (H), each carrying distinct biology, clinical behavior, and therapeutic vulnerability (Fig. 1).^20–22^ Naturally occurring canine invasive BC (InvBC) recapitulates this organization with striking fidelity: in a cohort of 56 dogs, the current concensus human classifier and unsupervised clustering both resolved two stable groups, 45 luminal and 11 basal (in contrast with 29 luminal and 27 basal with an older 60-gene classifier in the original publication), with similar immune-infiltration and clinical outcomes as seen in humans.^13, 14^ At the DNA level, however, the two species diverge sharply. While less common in human MIBC compared to NMIBC, *FGFR3* alterations are enriched in the luminal-papillary subtype of MIBC tumors, ^16, 23, 24^ while canine InvBC is dominated by the *BRAF* V595E mutation (present in 75% of dogs in the Sommer cohort), and this mutation is itself enriched in the canine luminal subtype.^13, 25^ Other recurrent alterations, including *TP53* and *TERT* promoter alterations, are important in MIBC;^16^ here we focus on *FGFR3* and *BRAF* because they both drive cancer through common MAPK signaling and BRAF is the only genetic data available for this Sommer cohort (Fig. 1A). The two driver lesions are non-orthologous as genes, but they are orthologous as pathway entries: both feed into RAS-RAF-MEK-ERK upstream of the proliferative transcriptional program that defines the luminal subtype.^26–29^ Thus *FGFR3* and *BRAF* variants in each species converge on a shared luminal and proliferative pathway program in both species.^6, 30^

**Figure 1.**
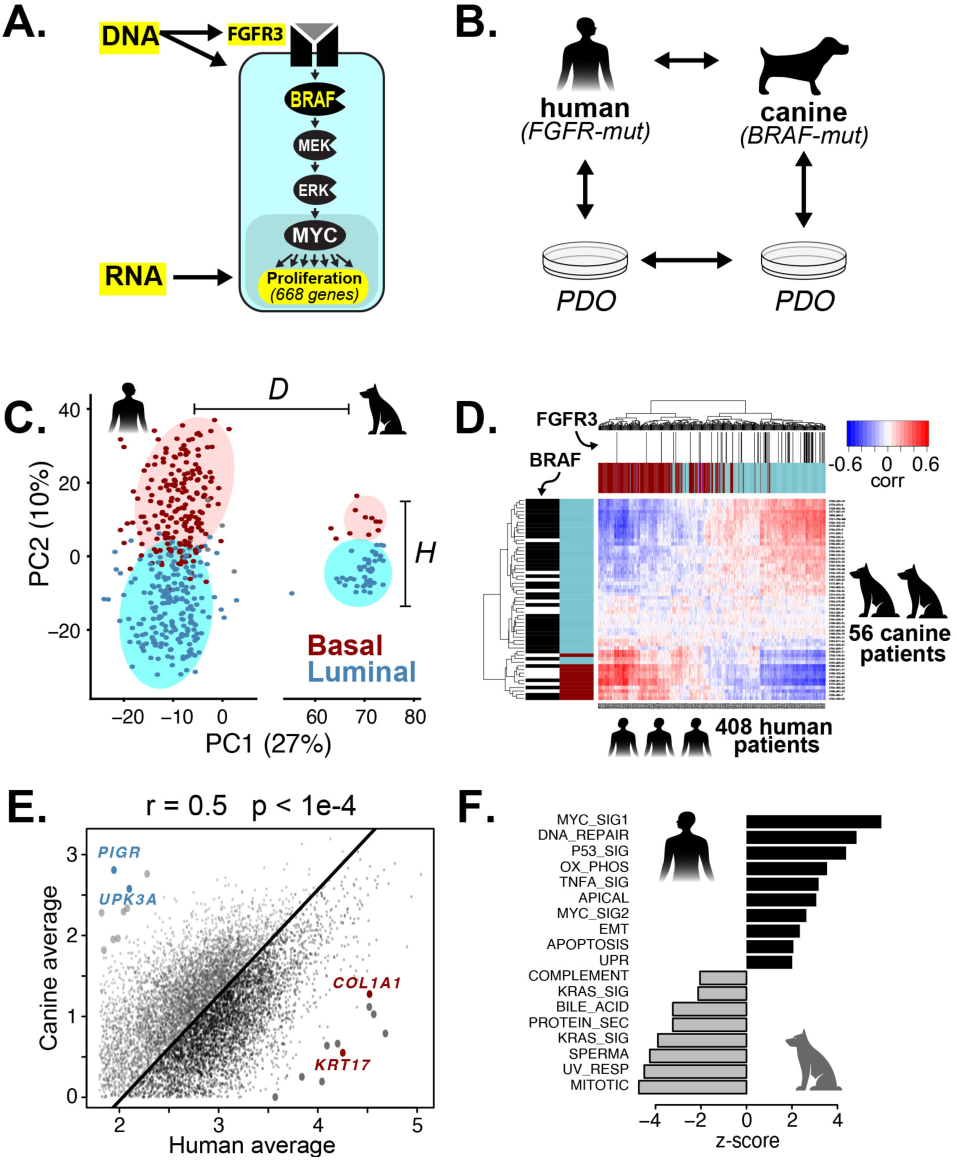
A cross-species framework for transcriptomic fidelity across species and platforms. **A.** DNA-level mutations cluster at pathway inputs, while RNA-level readouts report the downstream proliferative program licensed by the RAS-RAF-MEK-ERK cascade. **B.** The 2×2 fidelity loop. Human and canine patients and their patient-derived organoids define four nodes connected by four pairwise comparisons: each tests a distinct fidelity claim. **C.** Principal component analysis of human (n=408)^16^ and canine (n=56)^13^ InvBC patient transcriptomes. **D.** Cross-species correlation heatmap of human and canine patients over shared orthologous genes, annotated by FGFR3 and BRAF mutation status.^13,16^ **E.** Patient level averaging can reveal overall species-specific differences at the gene level. **F.** Pathway enrichment on species-averages reveals systematic differences between humans and canines.^13, 16^

While *in vivo* models are typically ideal, to increase throughput, further reduce costs, and address safety concerns like toxicity, the use of *in vitro* models is a critical necessity. With the advent of newer and more advanced *in vitro* models, such as spheroids, organoids, and organs-on-a-chip, increased fidelity with clinical complexity and heterogeneity is increasingly feasible. Functional precision oncology extends this paradigm by generating patient-derived organoids (PDOs), *ex vivo* models that retain many of the molecular characteristics of the tumors from which they were derived.^31, 32^ PDOs permit medium- to high-throughput therapeutic testing and controlled mechanistic experiments while preserving patient-specific biology, enabling both clinical applications such as drug repurposing and scientific studies of disease mechanisms. Their principal limitation is that they exist outside the native tissue environment.^33–35^ As a result, evaluating fidelity requires consideration of both species^36, 37^ and platform effects.^9, 17, 38^

Together, these considerations define a 2×2 fidelity loop spanning species and platform transitions (Fig. 1B). A useful model must preserve conserved biology (C), maintain clinically relevant heterogeneity (H), and distinguish these from systematic divergence (D) introduced by species or experimental platforms.

Despite this molecular parallelism, human and canine bladder cancer models have never been directly compared in a single integrated analysis. Here we close that gap by analyzing human and canine patient transcriptomes simultaneously to quantify inter-species divergence (D) and patient-level heterogeneity (H) on a common axis, and to map the correlation structure across luminal and basal molecular subtypes and clinical endpoints. We then extended the comparison to paired PDOs from both species, closing the 2×2 fidelity loop. Finally, we used single-cell RNA-seq^9, 18^ and spatial transcriptomic data^34^ to resolve cellular heterogeneity within these tumors and the degree to which it is conserved across species.

## RESULTS

### Species and platform differences separate cleanly from conserved luminal-basal biology (Fig 1)

#### Conceptual framework

We used a variance decomposition framework, described mathematically in Supplementary Note 1, to organize the analyses and figures around three sources of transcriptomic variation: divergence (D), conservation (C), and patient-heterogeneity (H). Divergence refers to systematic differences between species or experimental platforms, such as human versus canine tumors or tumors versus organoids. Conservation refers to biological patterns that remain recognizable across those differences, such as a shared luminal-basal axis. Heterogeneity refers to patient-to-patient variation within each group. PCA provides a visual summary of these relationships: in our analyses, PC1 often captured the largest species or platform difference, while PC2 captured the conserved luminal-basal structure. Patient/sample heterogeneity is reflected by the spread of individual samples along both axes.

#### Principal Component Analysis

We first applied this framework to bulk transcriptomes from human (n=408)^16^ and canine(n=56)^13^ InvBC tumors. Expression matrices were restricted to unambiguous 1-to-1 orthologs, combined, quantile normalized and analyzed by principal component analysis (Fig. 1C). Human and canine samples separated primarily along PC1, and global expression profiles differed significantly (PERMANOVA: F = 161.68, R^2^ = 0.262, p < 0.0001). Quantile normalization reduced platform-related distributional and scale differences but could not eliminate technical effects confounded with species. Accordingly, PCA was used to visualize the combined species and platform divergence rather than to correct it, because batch correction would artificially reduce the observed cross-cohort variation.

Although species and platform differences dominated PC1, both cohorts retained a luminal-basal axis along PC2, consistent with conserved tumor biology.^13^ Luminal-basal separation was strong in both humans (Cohen’s d = 2.84) and in dogs (Cohen’s d = 3.49) and was significantly associated with global gene variation (PERMANOVA: R^2^ = 0.077; p < 0.0001). After limma batch correction, this separation remained significant across species (PERMANOVA: R^2^ = 0.093, p < 0.0001) supporting the robustness of the conserved luminal-basal axis to platform related effects. Thus, despite dominant species- and platform-associated divergence (D), both cohorts retained a parallel luminal-basal structure representing conserved inter-patient heterogeneity (H; Fig. 1C).

#### Correlation Analysis reconciles DNA and RNA

To examine conserved structure among individual tumors, we computed pairwise correlations across all human and canine tumors (Fig. 1D). Within each species, tumors formed reciprocal luminal and basal blocks, indicating that molecular subtype is the major source of inter-patient variation. These blocks aligned the luminal subtypes with *FGFR3* mutations in human tumors and *BRAF* mutations in canine tumors, consistent with the association of both drivers with luminal disease.^6, 30^ Cross-species correlations were substantially weaker than within-species correlations (Fig. S2). The median correlation was 0.41 between human and canine tumors (IQR, 0.36–0.45), compared with 0.73 among human tumors (IQR, 0.67–0.78) and 0.89 among canine tumors (IQR, 0.86–0.91).

Species-level averaging of expression profiles nevertheless remained correlated across 9,569 orthologous genes (Fig. 1E; Pearson r = 0.50, 95% CI 0.49 to 0.52, n = 9,569 orthologous genes, p < 2.2 × 10^-16^). Although averaging removes patient-level variation, it preserves the cohort-level consequences of differing subtype composition. Canine-enriched outliers, including PIGR and UPK3A, marked luminal urothelial differentiation, whereas human-enriched outliers, including *COL1A1* and *KRT17*, reflected basal phenotypes.^39^ Thus, these deviations likely reflect the greater prevalence of luminal tumors in the canine cohort (∼80%) than in the human cohort (∼40%), rather than arbitrary species-specific differences. Gene set enrichment analysis (GSEA) applied to differential expression between canine-and human data revealed that human samples show statistically significant enrichment in proliferation pathways, epithelial-to-mesenchymal transition (EMT) and inflammatory pathways (Fig 1F).

Together, these analyses demonstrate a conserved transcriptional relationship between human and canine InvBC tumors but do not identify the underlying biological processes. We therefore next applied gene-signature analysis, which aggregates coordinated genes into more stable biological readouts (Fig. S3). We evaluated two complementary signature types: pathway signatures, which capture intracellular programs,^40^ and cell-type signatures, which resolve the cellular composition of each bulk tumor sample (Fig 2).^41^

**Figure 2.**
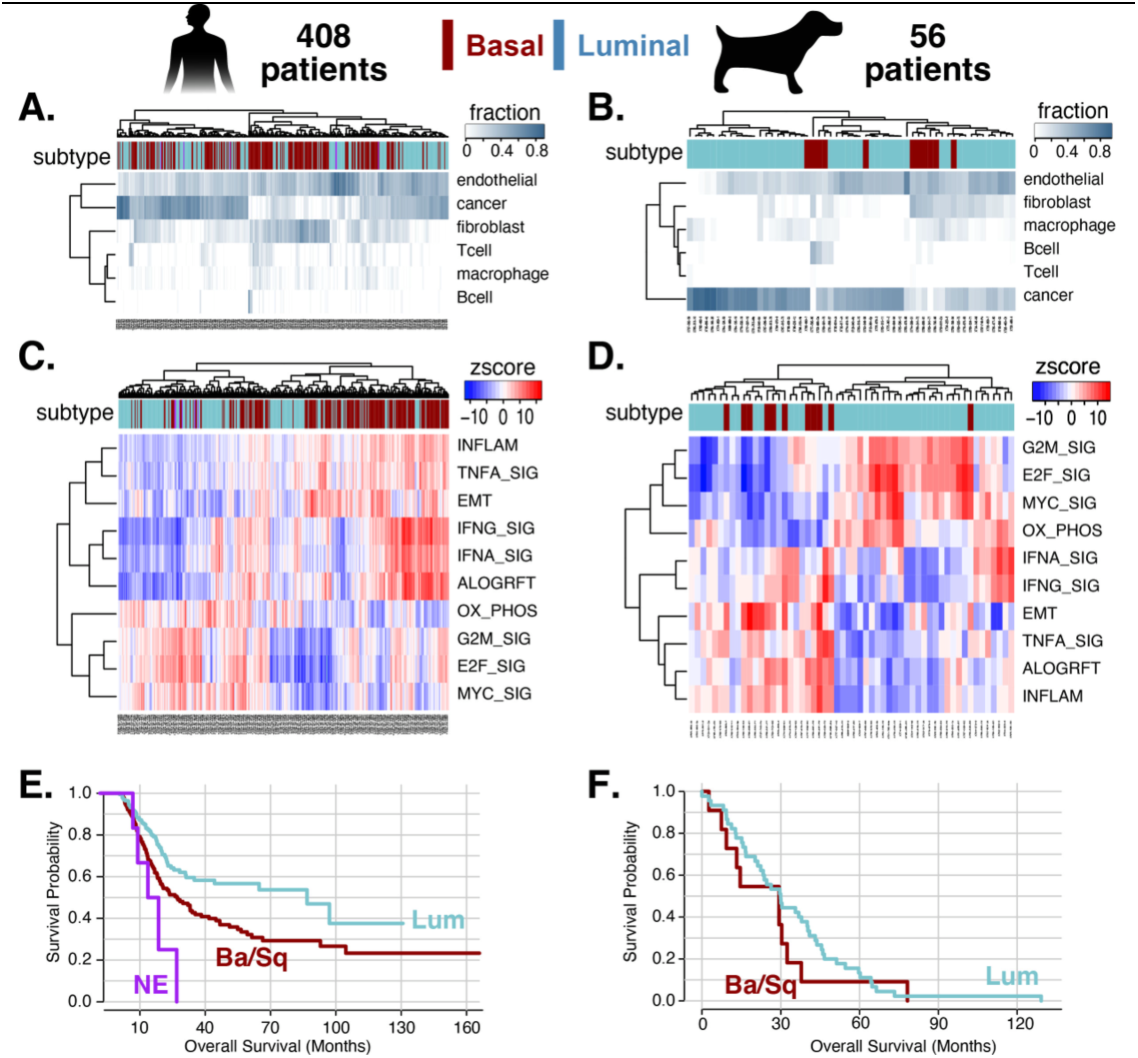
Cross-species comparison of human and canine InvBC across clinical, cellular and molecular features. All panels compare human (n=408) on the left and canine (n=56) InvBC cohorts on the right, with samples annotated by molecular subtype.^13, 16^ **(A-B)** Tumor microenvironment composition by cellular deconvolution (fraction per cell type) in human and canine tumors. **(C-D)** Pathway signature activity (z-score) in human and canine tumors, partitioning into proliferative (MYC, E2F, G2M) and inflammatory (IFNA, IFNG, TNFα, allograft rejection, EMT) modules. **(E-F)** Kaplan-Meier overall survival by subtype in human (E; luminal, basal/squamous, neuroendocrine) and canine (F; luminal, basal/squamous) cohorts.

### Clinical, pathway, and microenvironmental features are conserved across species

**CIBERSORT deconvolution** revealed a similar tumor-microenvironment structure in human and canine bladder cancers (Fig. 2A,B). ^13, 16^ In both species, tumors followed a subtype-associated gradient from epithelial-dominant to stromal- and immune-rich. Luminal tumors were enriched for epithelial cells, whereas basal tumors showed greater fibroblast, macrophage, T-cell, and B-cell infiltration; five of six inferred populations showed the same subtype association across species. Thus, molecular subtype reflects both tumor-cell-intrinsic programs and a conserved microenvironmental architecture, consistent with previous studies on smaller patient cohorts.^13, 34, 39^ Figures 4 and 5 examine this pattern further at single-cell and spatial resolution.

**Pathway analysis** identified the biological programs underlying this conserved structure (Fig. 2C,D). Of the 50 hallmark pathways evaluated in both datasets, 40 showed the same luminal-basal association in human and canine tumors (Fisher’s exact p = 1 × 10^-5^, OR = 22.8). In both species, pathways separated into reciprocal proliferative and inflammatory modules. Luminal tumors were enriched for MYC, E2F, and G2M programs, whereas basal tumors were enriched for interferon responses, TNFα signaling, allograft rejection, epithelial–mesenchymal transition, and other inflammatory programs. This pattern is consistent with the underlying genetics: FGFR3 alterations in human luminal tumors and BRAF mutations in canine luminal tumors both activate RAS-RAF-MEK-ERK signaling upstream of proliferative programs,^26–29^ whereas basal tumors exhibit an inflamed, mesenchymal, and stroma-rich phenotype.^20, 30, 34, 42, 43^ Thus, a conserved proliferation–inflammation axis organizes invasive bladder cancer in both species.^13^

#### Clinical Analysis

These conserved molecular features correlated with clinical outcome. In the human cohort, luminal tumors were associated with longer overall survival than basal tumors, with neuroendocrine tumors faring worst, reproducing the established prognostic ordering of MIBC subtypes (basal vs luminal HR = 1.72, 95% CI 1.26-2.34; log-rank p = 2 × 10^-4^) (Fig. 2E).^39, 44^ All 56 dogs experienced the survival event. Survival did not differ significantly between basal and **luminal** tumors by log-rank testing (p = 0.387). In a Cox proportional-hazards model, **basal** tumors had an HR of 1.35 relative to **luminal** tumors (95% CI, 0.69–2.65; p = 0.381). The proportional-hazards assumption was not violated (cox.zph, p = 0.406). However, the wide confidence interval indicates limited precision and does not exclude a clinically meaningful difference.^13^ Taken together, these results demonstrate that canine InvBC recapitulates human InvBC at at a broad phenotypic level.

### Tumor-to-organoid changes are conserved across species (Fig 3)

The analyses to this point compared human and canine tumors directly, establishing fidelity along the cross-species axis of the framework. The remaining axes involve patient-derived organoids (PDOs), the ex vivo models on which functional precision medicine relies. To close the loop, a model system must be credentialed not only across species but across the tumor-to-organoid transition (Fig. 3A-B).

**Figure 3.**
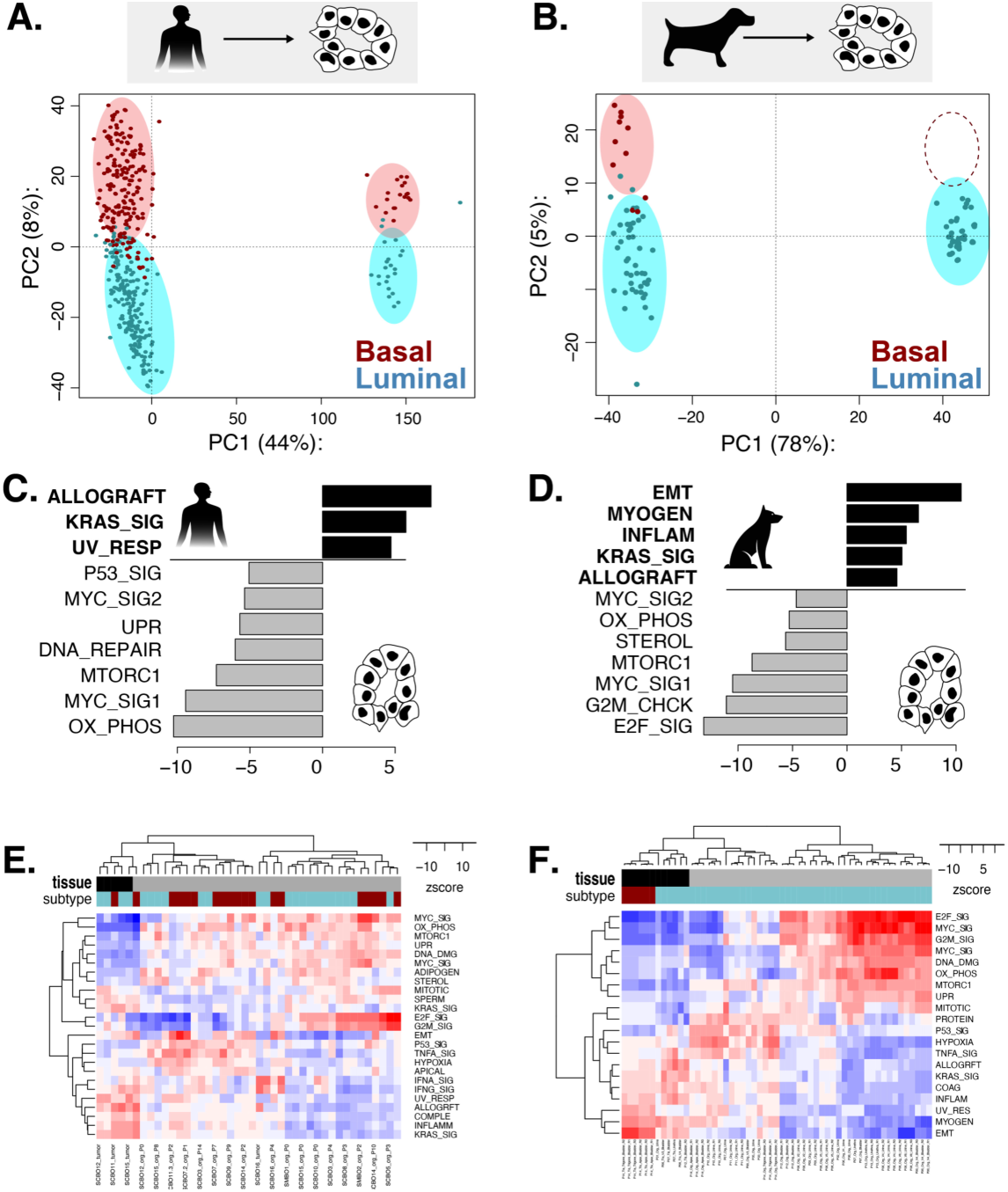
Cross-species comparison of human (left side) and canine (right side) InvUC tumors and InvUC-derived organoids. **(A-B)** Principal component analysis of tumors and organoids in the human and canine cohorts, colored by subtype (basal, red; luminal, cyan). PC1 separates tumors from organoids in both species; subtype organization is preserved along PC2 within the organoid compartment. **(C-D)** Top pathway signature activities (z-score) on average tumor and organoids, reveals conservation in organoid-culture bias. **(E-F)** Pathway signature activity (z-score) in human and canine tumors and organoids, annotated by tissue (tumor, black; organoid, grey) and subtype. Organoids are enriched for proliferative and metabolic programs; tumors retain inflammatory, EMT, and stromal programs.

We assembled four cohorts spanning the 2×2 design in Figure 1B. On the tissue side are the 408 human and 56 canine patient tumors analyzed above.^13, 16^ On the organoid side are 42 human bladder cancer PDO samples drawn from a published organoid biobank and 41 samples from 25 canine urothelial carcinoma-derived PDOs derived from tumors and/or urine from 14 patients.^17, 18^

**Principal component analysis** of tumors and organoids within each species revealed a consistent structure (Fig. 3). The dominant axis of variation (PC1) separated tumors from organoids, in both human (44% of variance) and canine (78%) cohorts. This separation was conserved across global expression profiles for both humans (PERMANOVA: F = 346.3, R2 = 0.439, p < 0.0001) and canine (PERMANOVA: F = 348.18, R^2^ = 0.780, p < 0.0001) patients.

The second component runs within each platform and separates ***luminal*** *(blue)* from ***basal (red)*** tumors for both species. In humans, molecular subtype remained significantly associated with global gene-expression profiles both before (PERMANOVA: F = 25.6, R^2^ = 0.055, p < 0.0001) and after limma batch-correction (PERMANOVA: F = 47.0, R^2^ = 0.096, p < 0.0001), supporting conservation of luminal and basal/squamous transcriptional structure across both organoid and tumor contexts. No canine PDOs were classified as basal (Fig 3B,F) and therefore subtype structure could not be evaluated.

**Pathway analysis** identified the biology underlying tumor-organoid divergence, and it was the same in human (Fig. 3 C) and canine (Fig. 3D). Organoids were enriched for proliferative and metabolic programs (i.e. MYC, E2F, and G2M targets, oxidative phosphorylation, mTORC1 signaling, and DNA-damage response) consistent with rapid growth in culture under defined media. Tumors retained the inflammatory, and stromal programs that depend on a microenvironment absent from organoid culture.

Taken together, the bulk RNA-seq comparison completes the four edges of the fidelity loop (Fig. 1B): human and canine tumors share conserved transcriptional programs, human and canine organoids share the same culture-induced shift characterized by increased proliferation and reduced enrichment of inflammatory genes. To better understand basal-luminal heterogenituy within tumors we next turned to single-cell resolution data across species and platforms (Fig 4).

**Figure 4.**
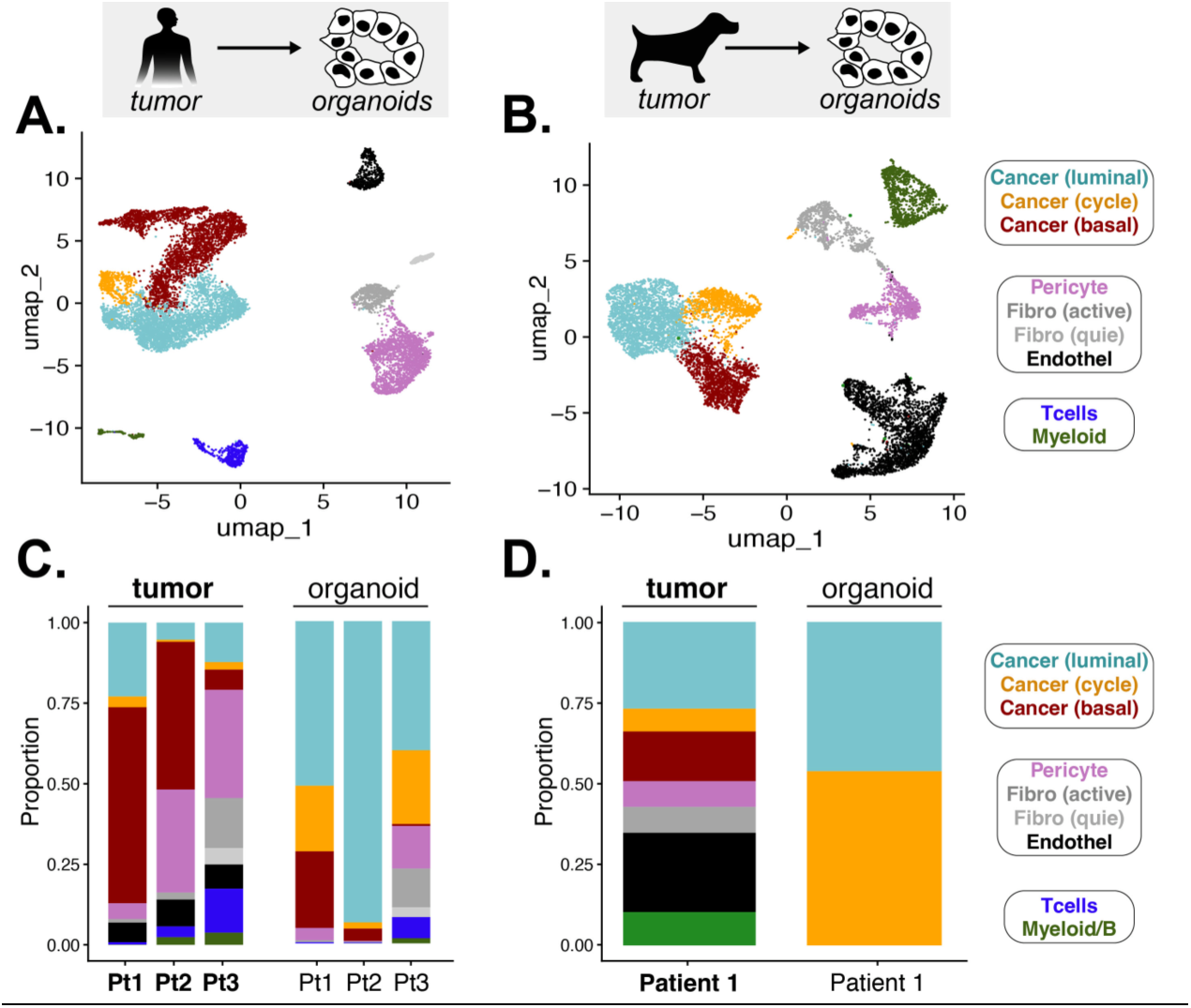
Cross-species comparison of human and canine tumors using single-cell RNA-seq. **(A-B)** UMAP visualizations of scRNA-seq from human and canine paired tumors and organoids, colored by cell type. Both species resolve the same compartments: three cancer states (luminal, cycling, and basal), stromal populations (pericytes, active and quiescent fibroblasts, endothelium), and immune populations (T cells, myeloid/B cells), occupying analogous regions of the UMAP. **(C-D)** Cell-type composition of paired tumor and organoid samples for three human patients (C) and one canine patient (D). Two changes occur together in the transition to culture: the stromal and immune compartments that make up a large fraction of every tumor are nearly absent from organoids, and within the surviving cancer compartment the balance shifts away from basal toward the luminal and cycling states. Both changes are also present in the one available canine pair (n = 1).

### PDO culture favors luminal and cycling epithelial states (Fig. 4)

Single-cell or single-nuclei RNA sequencing of paired tumors and organoids resolved the cellular composition underlying bulk profiling in the previous section.^9, 18^ The same compartments appeared in both species: a cancer compartment, stromal populations (quiescent and active fibroblasts, pericytes, endothelial cells), and immune populations (T cells, myeloid and B cells). The architecture of the two datasets was strikingly parallel, with cancer cells, stroma, and immune cells types and frequencies in human and canine alike. The cellular ecosystem of bladder cancer is therefore conserved across species not only as bulk transcriptional programs, but as discrete, identifiable cell types (Fig. 4A,B).

Within the cancer compartment, the data resolved three states rather than two. Alongside the expected luminal and basal clusters was a third, cycling population that is positioned in between them and leaned luminal. This corresponds to the state previously described as "intermediate", a transitional population bridging a basal-stem-like phenotype and the differentiated luminal and umbrella cells of normal urothelium.^34, 45^ Critically, individual tumors contained all three states simultaneously (Fig. 4C-D). Therefore the luminal-basal axis that bulk subtyping reduces to a single per-tumor label is, at single-cell resolution, a continuum of cancer-cell plasticity within each tumor. ^9, 34, 43, 45^

The paired composition analysis revealed two distinct changes occurring together in the transition from tumor to organoid (Fig. 4 C,D). First, the stromal and immune compartments, which made up a substantial fraction of every tumor, were almost entirely lost in culture: organoids retained essentially only epithelial cells.

Second, within the surviving cancer compartment, the balance of states shifted away from basal and toward the luminal and cycling states. Both changes were present in all three human pairs and consistent with the one available canine pair. While this interpretation is limited by the lack of basal canine tumor-organoid pairs, this decomposes the apparent "luminal shift" seen in the bulk data (Fig. 3) into its two real components: the loss of non-epithelial cells whose transcripts had diluted the luminal signal in tissue, and a genuine shift in epithelial state toward luminal identity. A recent proposal holds that the basal state is not cell-intrinsic but enforced by the surrounding niche and contacts between cancer cells and the immune and stromal populations near the basal membrane.^34^

### Human Spatial Transcriptomics Reveals Basal Cells are Enriched Along the Stromal Border (Fig. 5)

The mechanistic question raised by the previous section was, at its core, spatial: does the basal state occupy a defined position relative to the stromal and epithelial compartments? A single-cell-resolution spatial transcriptomic dataset of human MIBC, allows the prediction to be tested directly (Fig. 5).^34^ Mapping the same cell-state classification onto intact tissue resolves the identical compartments seen in dissociated single-cell data, the three cancer states and the full stromal and immune complement, but now with their positions preserved (Fig. 5). Basal cancer cells form a continuous rim along the stromal interface, lining the outer boundary of every epithelial nest where it abuts fibroblasts, endothelium, and infiltrating immune cells. Luminal cells fill the differentiated interior of each nest, away from the stroma. The cycling, intermediate state occupies a band between them, just interior to the basal rim.

**Figure 5.**
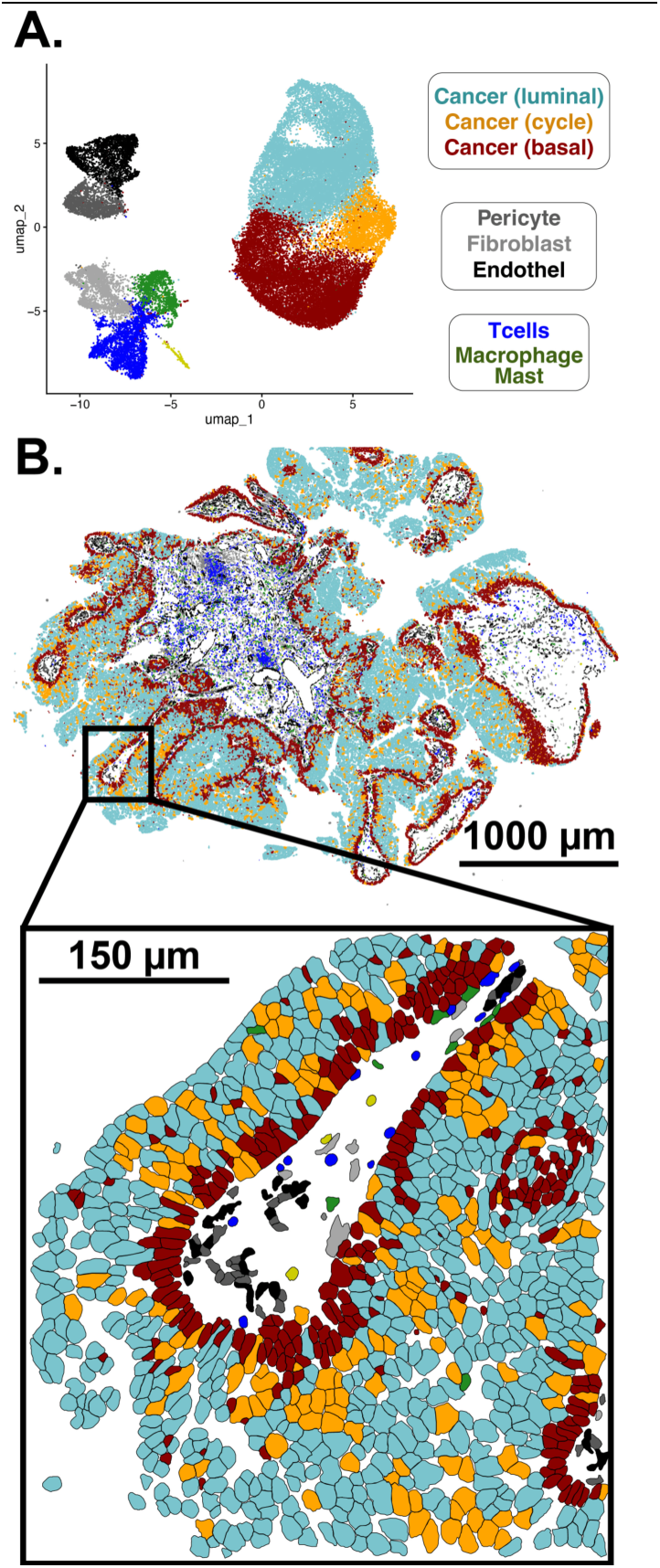
One representative human patient illustrating the spatial organization of MIBC cell states by single-cell-resolution spatial transcriptomics.^34^. **(A)** UMAP visualization of the spatial dataset colored by cell type, resolving the same compartments seen in dissociated single-cell data: three cancer states (luminal, cycling, basal), stromal populations (pericytes, fibroblasts, endothelium), and immune populations (T cells, macrophages, mast cells). **(B)** The same cell-state classification mapped back onto intact tissue, with each cell plotted at its spatial position. The arrangement is non-random: basal cancer cells (dark red) form a continuous rim along the outer boundary of each epithelial nest where it abuts the stromal and immune interstitium, luminal cells (cyan) fill the differentiated interior, and the cycling state (orange) occupies a band between them, just interior to the basal rim. This geometry recasts the luminal-basal axis as a differentiation gradient organized around the stromal boundary

This geometry reconciles the findings of the preceding sections into a single coherent picture. The luminal-basal axis that bulk subtyping treats as a per-tumor label is, spatially, a differentiation gradient organized around the stromal boundary: basal at the niche interface, intermediate in transit, luminal in the interior. The same gradient may explain the organoid result, since culture removes the stromal and immune partners that define the basal interface, and cancer cells deprived of that boundary relax toward the luminal interior state that now has nowhere to orient against. It also could explain why the cross-species conservation documented throughout holds at the level of programs, cell states, and now tissue architecture, while bulk subtype calls remain coarse: both species build the same spatially organized ecosystem, and bulk measurement averages across it.

## DISCUSSION

We asked three related questions: which features of human and canine InvBC bladder cancer are conserved, which diverge (e.g. barplots Fig 1-3), and whether clinically relevant patient heterogeneity is preserved in tumors and patient-derived organoids (PDOs). Although species and platform dominated the largest axis of transcriptomic variation, both species retained the same luminal-basal organization, proliferation-inflammation axis, and subtype-associated tumor-microenvironment structure. This shared biology arose through different genetic drivers (e.g. FGFR3 vs BRAF) that converge on similar downstream pathways. Thus, conservation is strongest at the level of cell states and biological programs.

Single-cell analysis refined the bulk subtype framework by showing that luminal, cycling, and basal cancer-cell states coexist along a shared differentiation continuum in both human and canine bladder cancer.^9, 18^ Comparing primary tumors with matched PDOs further showed that organoid culture alters the relative abundance of these conserved states, shifting cells toward a more luminal and proliferative phenotype. Thus, bulk subtype differences reflect changes in the composition of conserved cancer-cell states, while PDOs preserve only part of the heterogeneity present in primary tumors.

Human spatial data provided additional context for this tumor-organoid divergence.^34^ Basal cells localized near the stromal interface, luminal cells occupied the differentiated epithelial interior, and cycling cells were positioned between these regions. This organization suggests that interactions with the surrounding microenvironment help maintain the basal state and that their loss during organoid culture contributes to the luminal shift.^17^ Our contribution is therefore the integration of cross-species single-cell comparisons with tumor-PDO analyses, using human spatial data to provide a mechanistic explanation for the loss of cellular heterogeneity in culture. These findings complement prior spatial studies linking stromal architecture to clinical outcomes in human MIBC.^46^

Clinical outcomes were directionally consistent but less definitive than these molecular findings. Basal disease was associated with poorer survival in both species, but the canine result did not reach statistical significance and was sensitive to classifier choice.^13^ Cross-species comparison is further complicated by major differences in care. The most consequential is systemic therapy: human MIBC is commonly treated with neoadjuvant or adjuvant cisplatin-based chemotherapy, and increasingly with immune checkpoint inhibitors and antibody-drug conjugates,^47^ whereas dogs are typically managed with NSAIDs such as piroxicam, with or without cytotoxics such as mitoxantrone, vinblastine, and carboplatin.^48^ Local therapy differs as well: human patients may undergo radical cystectomy, including removal of the bladder and regional lymph nodes,^47^ whereas dogs rarely receive definitive local therapy and more often undergo partial tumor resection or medical therapy alone.^48^ Canine survival data are also subject to informative censoring: death is frequently elective euthanasia prompted by progressive symptoms, quality of life, prognosis, or financial constraints. The canine endpoint is therefore overall survival rather than cancer-specific survival, and these competing risks are not independent of disease burden. Thus, the canine data support a possible conserved prognostic relationship, but the systemic therapy gap and end-of-life practice make the magnitude of that relationship non-comparable across species.

### Limitations

This study has several limitations. First, the human and canine cohorts were generated on different sequencing platforms and mapped to different reference genomes (see data sources in methods below). Quantile normalization harmonizes the overall expression distributions but does not remove platform-specific structure, and that structure loads onto PC1. The between-species distance along PC1 therefore reflects a mix of biological divergence and technical batch effect and should not be read as a purely biological measure. The conserved luminal-basal axis on PC2 survives explicit batch correction and does not carry this caveat. Second, no canine organoids were classified as basal, so the tumor-to-organoid comparison in dogs could be evaluated only for luminal tumors, and whether canine basal tumors retain their subtype in culture remains untested. Third, the canine survival difference between subtypes was not significant in our analysis (HR 1.35, 95% CI 0.69 to 2.65, p = 0.38), whereas an earlier study reported a significant difference using a different molecular classifier (p = 0.0113). Because the shared canine cohort is small and every dog experienced the event, this disagreement most likely reflects sensitivity to classifier choice rather than a true absence of prognostic signal, but our data cannot settle which classifier is correct. Fourth, several of the cross-species instruments are calibrated on human data: the CIBERSORT reference and the DoRothEA regulons were both built from human samples and applied to canine transcriptomes through ortholog mapping. Prior benchmarking supports cross-species transfer of transcription-factor regulons,^49, 50^ but we cannot exclude that some of the apparent conservation reflects shared human-derived instrumentation rather than shared canine biology, and canine-specific references would be needed to separate the two. Fifth, canine-genetic data was limited to BRAF variants, preventing comprehensive analysis of variants between species. Finally, the single-cell tumor-to-organoid comparison rests on three human pairs and only one canine pair, so the canine result is a consistency check rather than an independent replication and will require additional animals to confirm.

### Future directions

Human spatial data showed that basal cancer cells are concentrated along the tumor–stroma boundary, which may explain why they are lost when tumors are grown as PDOs. Mapping canine tumors in the same way will show whether dogs share this tissue organization. Human and canine PDOs could also be grown with stromal and immune cells to test whether these supporting cells restore basal cancer states and their interactions. These experiments would clarify how tumors shift between basal and luminal states and how well canine models reflect human disease.

## METHODS

### Data Sources

Human muscle-invasive bladder cancer (MIBC) transcriptomic and clinical data were obtained from The Cancer Genome Atlas muscle invasive bladder cancer cohort (TCGA-MIBC; n = 408).^16^ Canine urothelial carcinoma transcriptomic data were obtained from a published cohort of 56 naturally occurring canine bladder cancers (GSE110661).^13^ Human bladder cancer patient-derived organoid (PDO) transcriptomic data were obtained from the bladder cancer organoid biobank reported by Lee et al. (GSE103990),^17^ while canine PDO data were obtained from a recently established canine urothelial carcinoma-derived organoid biobank comprising 52 mRNA profiles (11 tissue, 41 organoid) derived from tumor and/or urine samples producing 25 unique organoid lines from 14 dogs (GSE306810).^18^ Human and canine single-cell RNA sequencing datasets from tumors and matched organoid cultures were used to characterize cellular composition and lineage states across species (GSE217956, GSE306811). ^9, 18^ Spatial transcriptomic analyses utilized publicly available Xenium datasets from five human MIBC specimens to evaluate the spatial organization of epithelial, stromal, vascular, and immune cell populations (GSE326226). ^34^

All sample collections derived from canine patients at Iowa State University (IACUC-21-250) and the University of Georgia (A2023 10-002-A1) were used under approved IACUC protocols. All biospecimen collections obtained from Purdue University were conducted under an approved IACUC protocol (#1111000124), and tissue or urine samples were shipped to ISU or UGA.

### Clinical Data Analyses

#### Molecular Subtype Assignment and Integrated Transcriptomic Analyses

Human muscle- invasive bladder cancer (MIBC) transcriptomes were obtained from The Cancer Genome Atlas (TCGA-BLCA) cohort (n = 408),^16^ while canine InvBC transcriptomes were obtained from a published integrated canine bladder cancer cohort (n = 56).^13^ For cross-species analyses, expression matrices were restricted to unambiguous one-to-one human-canine orthologs identified using the orthogene R package, and quantile normalized. Human RNA-seq data were log-transformed and standardized prior to downstream analyses. Z-score normalized expression matrices were generated for analyses requiring direct comparison across samples and species. Quantile normalization harmonized the marginal expression distributions across the cohorts so that PCA and correlation were not driven by platform-level scale or depth; it does not remove the platform- and reference genome differences confounded with species. PCA was used precisely to visualize this combined species-and-platform effect rather than correct it, since preserving the magnitude of cross-cohort deviation, whether biological or technical, is itself informative (as any batch correction would coerce the data and obscure it). To confirm the conserved luminal-basal axis was not an artifact of this uncorrected structure, limma batch correction was applied and PERMANOVA on the corrected data confirmed that this luminal-basal separation was preserved across species.

Molecular subtypes were assigned using the Consensus Molecular Classification of Muscle-Invasive Bladder Cancer implemented through the consensusMIBC R package.^39, 51^ Tumors were initially classified into the six consensus subtypes (Luminal Papillary, Luminal Non-Specified, Luminal Unstable, Basal/Squamous, Stroma-rich, and Neuroendocrine-like). For cross-species comparisons, Luminal Papillary, Luminal Non-Specified, and Luminal Unstable tumors were collapsed into a single Luminal category, while Stroma-rich tumors were grouped with Basal/Squamous tumors.

To visualize global transcriptional relationships, principal component analysis (PCA) was performed on the quantile normalized human-canine expression matrix restricted to shared orthologous genes. Principal components were used to evaluate species-level divergence and subtype-associated transcriptional heterogeneity. Sample-to-sample similarity was further assessed using Pearson correlation analysis. Cross-species correlation heatmaps were generated using genes from a previously published canine MIBC molecular classifier, and correlation matrices were hierarchically clustered using average-linkage clustering. Human tumors were annotated with consensus molecular subtype and recurrent genomic alterations including FGFR3, TP53, RB1, and NFE2L2 mutations, while canine tumors were annotated with molecular subtype and BRAF mutation status as side bars on Figure 1D and S1B.

#### Cross-Species Conservation Analyses

To quantify transcriptomic conservation between human and canine bladder cancer, species-level average expression profiles were generated by averaging expression values across all tumors within each species. Cross-species concordance was assessed using Pearson correlation across shared orthologous genes. To identify genes exhibiting species-biased expression, orthogonal regression was performed on species-average expression profiles and orthogonal residuals were calculated for each gene. Genes with the largest positive or negative residuals were considered species-enriched, while genes with minimal residuals were considered highly conserved.

To place cross-species concordance in biological context, distributions of within-species and between-species correlations were compared (Fig S2). Pairwise Pearson correlations were calculated among human tumors, among canine tumors, and between all human-canine tumor pairs. Correlation distributions were used to evaluate whether cross-species conservation approached the level of variability observed among tumors within the same species.

#### Pathway, Transcription Factor, and Tumor Microenvironment Analyses

Pathway activity was inferred using the Virtual Inference of Protein-Activity by Enriched Regulon Analysis (VIPER) framework.^52^ Hallmark gene sets were obtained from the Molecular Signatures Database (MSigDB Hallmark collection) using the msigdbr package and converted into regulon objects for VIPER analysis.^40^ Sample-level pathway activity scores were calculated independently for human and canine cohorts. For visualization, Hallmark pathways were ranked by variance across samples within each cohort and the most variable pathways were displayed in heatmaps.

Transcription factor (TF) activity was inferred using the DoRothEA regulatory network resource and VIPER.^50, 52, 53^ Cross-species DoRothEA regulons with confidence levels A-C were used to estimate regulator activity from bulk transcriptomic profiles. TF activity scores were calculated from coordinated expression changes among downstream target genes and compared between species using the same conservation framework applied to gene-expression and pathway-level analyses. The human-derived regulons were applied to both species on shared orthologous genes, giving an identical instrument in each cohort; this transfer is supported by prior benchmarking showing that human TF footprints recover across mammalian models.^49, 50^

To assess conservation of higher-order biological programs, species-level Hallmark pathway activities and transcription factor activities were compared using orthologous pathway and regulator representations. Concordance was evaluated using species-average activity profiles and distributions of within-species versus between-species correlations (Fig S4).

Tumor microenvironment composition was estimated using the CIBERSORT algorithm based on ν-support vector regression.^41^ CIBERSORT was run in relative mode with 1000 permutations. Relative cell-type abundances were estimated from bulk transcriptomic profiles using an LM22-derived immune reference matrix. Resulting cellular fractions were used to compare immune and stromal composition across molecular subtypes and species. As above, the same reference was applied to both species on shared orthologous genes, prioritizing like-for-like cross-species comparisons over species-optimized deconvolution,

#### Survival Analyses

Overall survival analyses were performed using the survival and rms R packages. Kaplan-Meier survival curves were generated for molecular subtypes in both human and canine cohorts, and statistical significance was assessed using log-rank tests. Hazard ratios with 95% confidence intervals were estimated using Cox proportional hazards models, and the proportional-hazards assumption was assessed using scaled Schoenfeld residuals (cox.zph). Human survival analyses utilized TCGA clinical annotations,^16^ whereas canine analyses utilized matched clinical outcome data from the integrated canine cohort.^13^ Survival analyses were performed using the simplified molecular subtype assignments described above. The number of events and subjects at risk is reported for each cohort.

#### Statistical Analysis

All analyses were performed in R v4.5.2. Key package versions were Seurat 5.5.1, SeuratObject 5.4.0, harmony 2.0.5, viper 1.44.0, orthogene 1.16.1, and consensusMIBC 1.1.0, with the full environment pinned in the repository. Random seeds were fixed with set.seed(42) before Harmony integration, UMAP, and clustering so that embeddings and cluster assignments are reproducible. Principal component analysis was performed using prcomp after quantile normalization of shared gene-expression matrices, and the resulting components were used to visualize species and platform level structure. Species-level and platform level separation, and luminal versus basal separation within each species was evaluated by PERMANOVA using Euclidean distances with 9,999 permutations. Cross-species conservation of average gene or signature activity was assessed using Pearson correlation with 95% confidence intervals. Concordance of subtype-associated pathway and cell-state classifications between human and canine cohorts was evaluated using Fisher’s exact tests. Across the pathway-, transcription factor- and cell-type-level analyses, p-values were corrected for multiple comparisons using Benjamini-Hochberg procedure within each family of tests, and for confirmatory tests the Holm method was used. Adjusted p-values were reported. Overall survival was analyzed using Kaplan-Meier estimates, log-rank tests, and Cox proportional hazards models to estimate hazard ratios and 95% confidence intervals. The proportional hazards assumption was assessed using scaled Schoenfeld residuals and no significant violation was detected.

### Patient-derived organoid analyses

#### Patient-Derived Organoid Bulk RNA-seq Analyses

Human bladder cancer patient-derived organoid (PDO) bulk RNA-seq data were obtained from the published Lee et al. organoid cohort (GSE103990).^17^ Normalized count data were converted to reads per million (RPM), log10-transformed as log10(RPM + 1), and z-score normalized across genes where appropriate. Sample metadata were used to assign PDOs to their corresponding patient identifiers. Canine bladder cancer PDO RNA-seq data were obtained from an in-house canine organoid cohort that comprised 41 samples producing 25 unique organoid lines from derived from tumor and/or urine samples from 14 dogs.^18^ Expression values were analyzed as logTPM and z-score normalized matrices. Organoid samples were identified from sample annotations and separated from matched or associated tumor samples before downstream comparison.

Human and canine PDOs were molecularly subtyped using the consensusMIBC R package. Consensus classes were collapsed into simplified Luminal and Basal/Squamous categories using the same subtype definitions applied to the primary tumor cohorts. Hallmark pathway activity and transcription factor activity were inferred in PDO datasets using the same VIPER-based MSigDB Hallmark and DoRothEA workflows described above.

To compare primary tumors and PDOs, expression matrices were restricted to genes shared between the tumor and organoid datasets within each species. PERMANOVA and PCA was performed on combined tumor-organoid expression matrices using prcomp. These analyses were used to evaluate the extent to which PDO culture introduces a global transcriptional shift while preserving luminal-basal subtype structure. For Figure 3 heatmaps, Hallmark and transcription factor activity matrices were ranked by variance across PDO samples, and the most variable signatures were visualized with subtype annotations.

#### Single-Cell RNA-seq Analyses

Human and canine single-cell or single-nucleus RNA-seq datasets were analyzed using Seurat. Human data included matched bladder tumor and patient-derived organoid samples. Raw human count matrices were loaded from 10x Genomics HDF5 files and converted into Seurat objects using CreateSeuratObject, retaining genes detected in at least three cells and cells with at least 200 detected features. Sample metadata were added to each cell barcode and used to annotate patient identity, sample identity, and platform, defined as tumor or organoid.

Quality control was performed by calculating the number of detected genes, total RNA counts, and mitochondrial transcript fraction for each cell. Human cells were retained if they had at least 200 detected features and no more than 12% mitochondrial reads. Data were normalized using NormalizeData, highly variable features were selected using FindVariableFeatures, and expression values were scaled using ScaleData. Principal component analysis was performed using variable features.

To account for patient and sample effects, human tumor and organoid single-cell data were integrated using Harmony, with sample identity used as the batch variable. Neighbor graphs, clustering, and UMAP embeddings were then generated from Harmony-corrected dimensions. Clustering resolution was adjusted to identify both broad cell classes and finer epithelial and stromal subpopulations.

Cell-type annotation was performed by identifying cluster-enriched marker genes using FindAllMarkers with positive markers, a minimum expression fraction of 25%, and a log2 fold-change threshold of 0.25. Clusters were manually assigned to biological cell types using canonical marker genes. Luminal epithelial cells were identified by urothelial differentiation markers including UPK1A, UPK1B, UPK2, UPK3A, UPK3B, PSCA, and TFF family genes. Basal/squamous epithelial cells were identified by markers including KRT5, KRT17, S100A2, LAMB3, LAMC2, SERPINB5, and TP63-associated programs. Cycling epithelial cells were identified by proliferation markers including MKI67, TOP2A, CDK1, AURKB, BIRC5, and CCNB2. Stromal, vascular, and immune populations were annotated using markers of fibroblasts, pericytes, endothelial cells, T cells, myeloid cells, and B cells.

Canine single-nucleus RNA-seq analyses were performed using the same Seurat-based framework. Canine single-nucleus cells were retained with at least 200 detected features and no more than 5% mitochondrial reads. For analyses comparing canine tumor and organoid samples, tumor and organoid objects were merged, normalized, scaled, and integrated using Harmony with platform as the integration variable. Because canine organoid samples were strongly epithelial and did not contain the full stromal and immune diversity present in tumor tissue, tumor-only canine analyses were also performed to generate a clearer census of the *in vivo* tumor microenvironment. Canine tumor clusters were annotated using marker genes defining luminal epithelial, basal epithelial, cycling epithelial, endothelial, myeloid, fibroblast, and pericyte/smooth muscle populations.

Final cell-type labels were collapsed into harmonized categories to enable comparison across species and platforms. Human categories included luminal epithelial, basal epithelial, cycling epithelial, active fibroblast, quiescent fibroblast, endothelial, pericyte, T cell, and mixed myeloid/B-cell populations. Canine categories included luminal epithelial, basal epithelial, cycling epithelial, fibroblast, endothelial, pericyte, myeloid, and T-cell populations. UMAP plots were generated using final cell-type annotations (Fig 4A-B), and cell-type composition was summarized by calculating the proportional abundance of each annotated cell class within each sample, patient, or platform (Fig 4C-D). These proportional cell-type censuses were visualized as stacked bar plots to compare tumor and organoid cellular composition within and across species.

#### Xenium Spatial Transcriptomics Analysis

Human bladder cancer Xenium spatial transcriptomics data were analyzed using Seurat. Xenium cell-feature matrices were loaded from 10x Genomics HDF5 files using Read10X_h5, and the gene-expression matrix was used to create a Seurat object with the Xenium assay. Cell-level metadata, including centroid coordinates, were loaded from the corresponding cells.csv.gz file and added to the Seurat object. Cell identifiers were prefixed with the sample identifier to ensure unique cell names across downstream spatial and polygon-based analyses.

Quality control was performed using total Xenium transcript counts and detected feature counts per cell. Cells with low transcript counts were removed, and spatial distributions of transcript depth were visualized using centroid coordinates to assess regional technical variation. Xenium data were normalized using NormalizeData, variable features were identified with FindVariableFeatures, expression values were scaled with ScaleData, and principal component analysis was performed using RunPCA. Clustering was performed using FindNeighbors and FindClusters, followed by UMAP visualization using RunUMAP.

Cell-type annotations were assigned by identifying cluster-enriched marker genes with FindAllMarkers and manually mapping clusters to biological cell states. Luminal epithelial tumor cells were identified by urothelial differentiation markers including UPK1A, UPK2, UPK3A, ERBB2, ERBB3, MUC1, and PTK6. Basal epithelial tumor cells were identified by TP63, ITGB4, COL17A1, CLDN1, BCAM, CDH3, and related basal/stem-like markers. Cycling epithelial cells were identified by TOP2A, TYMS, CCNB1, CDC20, FOXM1, and CDK1. Stromal and immune populations were annotated using endothelial, fibroblast, pericyte, T-cell, macrophage, and mast-cell markers.

Annotated Xenium cells were visualized in both UMAP and tissue-coordinate space (Fig 5). Spatial plots were generated using cell polygons, fixed aspect ratios, reversed y-axis coordinates, and manually defined cell-type color palettes. To generate full-resolution spatial cell maps, cell-boundary coordinates were loaded from Xenium boundary parquet files using arrow, grouped by cell identifier, converted into closed polygons, and stored as sf polygon objects. Cell-type labels from the Seurat object were matched to polygon cell identifiers, allowing each segmented cell boundary to be rendered by annotated cell type using ggplot2 and sf. Large-scale spatial maps were exported as high-resolution images and converted to pyramidal TIFF/OME-TIFF format for visualization in QuPath when needed.^54^

## Data Availability

All data processing, statistical analyses and figure scripts are available via GitHub (https://github.com/fingolfn/POH_mibc). All datasets used are currently publicly available through Gene Expression Omnibus accession numbers listed above.

## REFERENCES

(1) Alghamdi, S. M.; Schofield, P. N.; Hoehndorf, R. Contribution of model organism phenotypes to the computational identification of human disease genes. Dis Model Mech 2022, 15 (7). DOI: 10.1242/dmm.049441.

(2) Hedges, S. B. The origin and evolution of model organisms. Nat Rev Genet 2002, 3 (11), 838–849. DOI: 10.1038/nrg929.

(3) Knapp, D. W.; Dhawan, D.; Ramos-Vara, J. A.; Ratliff, T. L.; Cresswell, G. M.; Utturkar, S.; Sommer, B. C.; Fulkerson, C. M.; Hahn, N. M. Naturally-Occurring Invasive Urothelial Carcinoma in Dogs, a Unique Model to Drive Advances in Managing Muscle Invasive Bladder Cancer in Humans. Front Oncol 2019, *9*, 1493. DOI: 10.3389/fonc.2019.01493.

(4) LeBlanc, A. K.; Breen, M.; Choyke, P.; Dewhirst, M.; Fan, T. M.; Gustafson, D. L.; Helman, L. J.; Kastan, M. B.; Knapp, D. W.; Levin, W. J.;, et al. Perspectives from man’s best friend: National Academy of Medicine’s Workshop on Comparative Oncology. Sci Transl Med 2016, 8 (324), 324ps325. DOI: 10.1126/scitranslmed.aaf0746.

(5) Ineichen, B. V.; Furrer, E.; Gruninger, S. L.; Zurrer, W. E.; Macleod, M. R. Analysis of animal-to-human translation shows that only 5% of animal-tested therapeutic interventions obtain regulatory approval for human applications. PLoS Biol 2024, 22 (6), e3002667. DOI: 10.1371/journal.pbio.3002667.

(6) Doncheva, N. T.; Palasca, O.; Yarani, R.; Litman, T.; Anthon, C.; Groenen, M. A. M.; Stadler, P. F.; Pociot, F.; Jensen, L. J.; Gorodkin, J. Human pathways in animal models: possibilities and limitations. Nucleic Acids Res 2021, 49 (4), 1859–1871. DOI: 10.1093/nar/gkab012.

(7) Beinse, G.; Tellier, V.; Charvet, V.; Deutsch, E.; Borget, I.; Massard, C.; Hollebecque, A.; Verlingue, L. Prediction of Drug Approval After Phase I Clinical Trials in Oncology: RESOLVED2. JCO Clin Cancer Inform 2019, 3, 1–10. DOI: 10.1200/CCI.19.00023.

(8) Ouimet, C.; Fodor, B.; Del Paggio, J. C.; Kimmelman, J. Proportion of patients in phase 2 oncology trials receiving treatments that are ultimately approved. J Natl Cancer Inst 2025, 117 (5), 1056– 1063. DOI: 10.1093/jnci/djaf013 From NLM Medline.

(9) Minoli, M.; Cantore, T.; Hanhart, D.; Kiener, M.; Fedrizzi, T.; La Manna, F.; Karkampouna, S.; Chouvardas, P.; Genitsch, V.; Rodriguez-Calero, A.;, et al. Bladder cancer organoids as a functional system to model different disease stages and therapy response. Nat Commun 2023, 14 (1), 2214. DOI: 10.1038/s41467-023-37696-2.

(10) Mullenders, J.; de Jongh, E.; Brousali, A.; Roosen, M.; Blom, J. P. A.; Begthel, H.; Korving, J.; Jonges, T.; Kranenburg, O.; Meijer, R.;, et al. Mouse and human urothelial cancer organoids: A tool for bladder cancer research. Proc Natl Acad Sci U S A 2019, 116 (10), 4567–4574. DOI: 10.1073/pnas.1803595116.

(11) Schneider, B.; Balbas-Martinez, V.; Jergens, A. E.; Troconiz, I. F.; Allenspach, K.; Mochel, J. P. Model-Based Reverse Translation Between Veterinary and Human Medicine: The One Health Initiative. CPT Pharmacometrics Syst Pharmacol 2018, 7 (2), 65–68. DOI: 10.1002/psp4.12262.

(12) LeBlanc, A. K.; Mazcko, C. N. Improving human cancer therapy through the evaluation of pet dogs. Nat Rev Cancer 2020, 20 (12), 727–742. DOI: 10.1038/s41568-020-0297-3.

(13) Sommer, B. C.; Dhawan, D.; Ruple, A.; Ramos-Vara, J. A.; Hahn, N. M.; Utturkar, S. M.; Ostrander, E. A.; Parker, H. G.; Fulkerson, C. M.; Childress, M. O.;, et al. Basal and Luminal Molecular Subtypes in Naturally-Occurring Canine Urothelial Carcinoma are Associated with Tumor Immune Signatures and Dog Breed. Bladder Cancer 2021, 7 (3), 317–333. DOI: 10.3233/BLC-201523.

(14) Dhawan, D.; Hahn, N. M.; Ramos-Vara, J. A.; Knapp, D. W. Naturally-occurring canine invasive urothelial carcinoma harbors luminal and basal transcriptional subtypes found in human muscle invasive bladder cancer. PLoS Genet 2018, 14 (8), e1007571. DOI: 10.1371/journal.pgen.1007571.

(15) Rossman, P.; Zabka, T. S.; Ruple, A.; Tuerck, D.; Ramos-Vara, J. A.; Liu, L.; Mohallem, R.; Merchant, M.; Franco, J.; Fulkerson, C. M.;, et al. Phase I/II Trial of Vemurafenib in Dogs with Naturally Occurring, BRAF-mutated Urothelial Carcinoma. Mol Cancer Ther 2021, 20 (11), 2177–2188. DOI: 10.1158/1535-7163.MCT-20-0893.

(16) Robertson, A. G.; Kim, J.; Al-Ahmadie, H.; Bellmunt, J.; Guo, G.; Cherniack, A. D.; Hinoue, T.; Laird, P. W.; Hoadley, K. A.; Akbani, R.;, et al. Comprehensive Molecular Characterization of Muscle-Invasive Bladder Cancer. Cell 2017, 171 (3), 540–556 e525. DOI: 10.1016/j.cell.2017.09.007.

(17) Lee, S. H.; Hu, W.; Matulay, J. T.; Silva, M. V.; Owczarek, T. B.; Kim, K.; Chua, C. W.; Barlow, L. J.; Kandoth, C.; Williams, A. B.;, et al. Tumor Evolution and Drug Response in Patient-Derived Organoid Models of Bladder Cancer. Cell 2018, 173 (2), 515–528 e517. DOI: 10.1016/j.cell.2018.03.017.

(18) Zdyrski, C.; Pawlak, A.; Nicholson, H. F.; Corbett, M. P.; Catucci, M.; Cheville, J.; Cho, H.; Melvin, B. J.; Peng, J.; Saba, C.;, et al. Establishment of a canine urothelial carcinoma-derived organoid biobank: A platform for comparative and translational research. Clin Transl Med 2026, *16* (4), e70645. DOI: 10.1002/ctm2.70645.

(19) Patel, V. G.; Oh, W. K.; Galsky, M. D. Treatment of muscle-invasive and advanced bladder cancer in 2020. CA Cancer J Clin 2020, 70 (5), 404–423. DOI: 10.3322/caac.21631.

(20) Choi, W.; Porten, S.; Kim, S.; Willis, D.; Plimack, E. R.; Hoffman-Censits, J.; Roth, B.; Cheng, T.; Tran, M.; Lee, I. L.;, et al. Identification of distinct basal and luminal subtypes of muscle-invasive bladder cancer with different sensitivities to frontline chemotherapy. Cancer Cell 2014, 25 (2), 152–165. DOI: 10.1016/j.ccr.2014.01.009.

(21) Dadhania, V.; Zhang, M.; Zhang, L.; Bondaruk, J.; Majewski, T.; Siefker-Radtke, A.; Guo, C. C.; Dinney, C.; Cogdell, D. E.; Zhang, S.;, et al. Meta-Analysis of the Luminal and Basal Subtypes of Bladder Cancer and the Identification of Signature Immunohistochemical Markers for Clinical Use. EBioMedicine 2016, 12, 105–117. DOI: 10.1016/j.ebiom.2016.08.036.

(22) Yoshida, T.; Kates, M.; Fujita, K.; Bivalacqua, T. J.; McConkey, D. J. Predictive biomarkers for drug response in bladder cancer. Int J Urol 2019, 26 (11), 1044–1053. DOI: 10.1111/iju.14082.

(23) Kompier, L. C.; Lurkin, I.; van der Aa, M. N. M.; van Rhijn, B. W. G.; van der Kwast, T. H.; Zwarthoff, E. C. FGFR3, HRAS, KRAS, NRAS and PIK3CA Mutations in Bladder Cancer and Their Potential as Biomarkers for Surveillance and Therapy. Plos One 2010, *5* (11). DOI: ARTN e13821 10.1371/journal.pone.0013821.

(24) Pouessel, D.; Neuzillet, Y.; Mertens, L. S.; van der Heijden, M. S.; de Jong, J.; Sanders, J.; Peters, D.; Leroy, K.; Manceau, A.; Maille, P.;, et al. Tumor heterogeneity of fibroblast growth factor receptor 3 (FGFR3) mutations in invasive bladder cancer: implications for perioperative anti-FGFR3 treatment. Ann Oncol 2016, 27 (7), 1311–1316. DOI: 10.1093/annonc/mdw170.

(25) Decker, B.; Parker, H. G.; Dhawan, D.; Kwon, E. M.; Karlins, E.; Davis, B. W.; Ramos-Vara, J. A.; Bonney, P. L.; McNiel, E. A.; Knapp, D. W.;, et al. Homologous Mutation to Human BRAF V600E Is Common in Naturally Occurring Canine Bladder Cancer-Evidence for a Relevant Model System and Urine-Based Diagnostic Test. Mol Cancer Res 2015, 13 (6), 993–1002. DOI: 10.1158/1541-7786.Mcr-14-0689.

(26) Cronise, K. E.; Hernandez, B. G.; Gustafson, D. L.; Duval, D. L. Identifying the ErbB/MAPK Signaling Cascade as a Therapeutic Target in Canine Bladder Cancer. Mol Pharmacol 2019, *96* (1), 36–46. DOI: 10.1124/mol.119.115808.

(27) Mahe, M.; Dufour, F.; Neyret-Kahn, H.; Moreno-Vega, A.; Beraud, C.; Shi, M.; Hamaidi, I.; Sanchez-Quiles, V.; Krucker, C.; Dorland-Galliot, M.;, et al. An FGFR3/MYC positive feedback loop provides new opportunities for targeted therapies in bladder cancers. EMBO Mol Med 2018, 10 (4). DOI: 10.15252/emmm.201708163.

(28) Tan, T. Z.; Rouanne, M.; Tan, K. T.; Huang, R. Y.; Thiery, J. P. Molecular Subtypes of Urothelial Bladder Cancer: Results from a Meta-cohort Analysis of 2411 Tumors. Eur Urol 2019, 75 (3), 423–432. DOI: 10.1016/j.eururo.2018.08.027.

(29) Ascione, C. M.; Napolitano, F.; Esposito, D.; Servetto, A.; Belli, S.; Santaniello, A.; Scagliarini, S.; Crocetto, F.; Bianco, R.; Formisano, L. Role of FGFR3 in bladder cancer: Treatment landscape and future challenges. Cancer Treat Rev 2023, 115, 102530. DOI: 10.1016/j.ctrv.2023.102530.

(30) Hovelson, D. H.; Udager, A. M.; McDaniel, A. S.; Grivas, P.; Palmbos, P.; Tamura, S.; Lazo de la Vega, L.; Palapattu, G.; Veeneman, B.; El-Sawy, L.;, et al. Targeted DNA and RNA Sequencing of Paired Urothelial and Squamous Bladder Cancers Reveals Discordant Genomic and Transcriptomic Events and Unique Therapeutic Implications. Eur Urol 2018, 74 (6), 741–753. DOI: 10.1016/j.eururo.2018.06.047.

(31) Letai, A. Functional Precision Medicine: Putting Drugs on Patient Cancer Cells and Seeing What Happens. Cancer Discov 2022, 12 (2), 290–292. DOI: 10.1158/2159-8290.CD-21-1498.

(32) Letai, A.; Bhola, P.; Welm, A. L. Functional precision oncology: Testing tumors with drugs to identify vulnerabilities and novel combinations. Cancer Cell 2022, 40 (1), 26–35. DOI: 10.1016/j.ccell.2021.12.004.

(33) Wang, L.; Izadmehr, S.; Sfakianos, J. P.; Tran, M.; Beaumont, K. G.; Brody, R.; Cordon-Cardo, C.; Horowitz, A.; Sebra, R.; Oh, W. K.;, et al. Single-cell transcriptomic-informed deconvolution of bulk data identifies immune checkpoint blockade resistance in urothelial cancer. iScience 2024, 27 (6), 109928. DOI: 10.1016/j.isci.2024.109928.

(34) Yu, K.; Chen, J.; Chu, Y. Y.; Nair, S.; Crupi, E.; Hasanov, E.; Mei, Y.; Han, X.; Liu, Y.; Liu, Y.;, et al. A Spatial Atlas of Muscle-Invasive Bladder Cancer Reveals Lineage-Specific Vulnerabilities and Immune Architecture. Cancer Discov 2026. DOI: 10.1158/2159-8290.CD-26-0099.

(35) Lin, J.; Jiang, S.; Chen, B.; Du, Y.; Qin, C.; Song, Y.; Peng, Y.; Ding, M.; Wu, J.; Lin, Y.;, et al. Tertiary Lymphoid Structures are Linked to Enhanced Antitumor Immunity and Better Prognosis in Muscle-Invasive Bladder Cancer. Adv Sci (Weinh) 2025, 12 (7), e2410998. DOI: 10.1002/advs.202410998.

(36) Dhawan, D.; Paoloni, M.; Shukradas, S.; Choudhury, D. R.; Craig, B. A.; Ramos-Vara, J. A.; Hahn, N.; Bonney, P. L.; Khanna, C.; Knapp, D. W. Comparative Gene Expression Analyses Identify Luminal and Basal Subtypes of Canine Invasive Urothelial Carcinoma That Mimic Patterns in Human Invasive Bladder Cancer. Plos One 2015, 10 (9). DOI: ARTN e0136688 10.1371/journal.pone.0136688.

(37) Ramsey, S. A.; Xu, T. J.; Goodall, C.; Rhodes, A. C.; Kashyap, A.; He, J.; Bracha, S. Cross-species analysis of the canine and human bladder cancer transcriptome and exome. Gene Chromosome Canc 2017, *56* (4), 328–343. DOI: 10.1002/gcc.22441.

(38) Elbadawy, M.; Usui, T.; Mori, T.; Tsunedomi, R.; Hazama, S.; Nabeta, R.; Uchide, T.; Fukushima, R.; Yoshida, T.; Shibutani, M.;, et al. Establishment of a novel experimental model for muscle-invasive bladder cancer using a dog bladder cancer organoid culture. Cancer Sci 2019, 110 (9), 2806–2821. DOI: 10.1111/cas.14118.

(39) Kamoun, A.; de Reynies, A.; Allory, Y.; Sjodahl, G.; Robertson, A. G.; Seiler, R.; Hoadley, K. A.; Groeneveld, C. S.; Al-Ahmadie, H.; Choi, W.;, et al. A Consensus Molecular Classification of Muscle-invasive Bladder Cancer. Eur Urol 2020, 77 (4), 420–433. DOI: 10.1016/j.eururo.2019.09.006.

(40) Liberzon, A.; Birger, C.; Thorvaldsdottir, H.; Ghandi, M.; Mesirov, J. P.; Tamayo, P. The Molecular Signatures Database (MSigDB) hallmark gene set collection. Cell Syst 2015, 1 (6), 417–425. DOI: 10.1016/j.cels.2015.12.004.

(41) Chen, B.; Khodadoust, M. S.; Liu, C. L.; Newman, A. M.; Alizadeh, A. A. Profiling Tumor Infiltrating Immune Cells with CIBERSORT. Methods Mol Biol 2018, 1711, 243–259. DOI: 10.1007/978-1-4939-7493-1_12.

(42) Seiler, R.; Ashab, H. A. D.; Erho, N.; van Rhijn, B. W. G.; Winters, B.; Douglas, J.; Van Kessel, K. E.; Fransen van de Putte, E. E.; Sommerlad, M.; Wang, N. Q.;, et al. Impact of Molecular Subtypes in Muscle-invasive Bladder Cancer on Predicting Response and Survival after Neoadjuvant Chemotherapy. Eur Urol 2017, 72 (4), 544–554. DOI: 10.1016/j.eururo.2017.03.030.

(43) Warrick, J. I.; Sjodahl, G.; Kaag, M.; Raman, J. D.; Merrill, S.; Shuman, L.; Chen, G.; Walter, V.; DeGraff, D. J. Intratumoral Heterogeneity of Bladder Cancer by Molecular Subtypes and Histologic Variants. Eur Urol 2019, 75 (1), 18–22. DOI: 10.1016/j.eururo.2018.09.003.

(44) Lerner, S. P.; McConkey, D. J.; Hoadley, K. A.; Chan, K. S.; Kim, W. Y.; Radvanyi, F.; Hoglund, M.; Real, F. X. Bladder Cancer Molecular Taxonomy: Summary from a Consensus Meeting. Bladder Cancer 2016, 2 (1), 37–47. DOI: 10.3233/BLC-150037.

(45) Tran, M. A.; Youssef, D.; Shroff, S.; Chowhan, D.; Beaumont, K. G.; Sebra, R.; Mehrazin, R.; Wiklund, P.; Lin, J. J.; Horowitz, A.;, et al. Urine scRNAseq reveals new insights into the bladder tumor immune microenvironment. J Exp Med 2024, 221 (8). DOI: 10.1084/jem.20240045.

(46) Feng, C.; Wang, Y.; Song, W.; Liu, T.; Mo, H.; Liu, H.; Wu, S.; Qin, Z.; Wang, Z.; Tao, Y.;, et al. Spatially-resolved analyses of muscle invasive bladder cancer microenvironment unveil a distinct fibroblast cluster associated with prognosis. Front Immunol 2024, 15, 1522582. DOI: 10.3389/fimmu.2024.1522582 From NLM Medline.

47. DeVita, V. T.; Lawrence, T. S.; Rosenberg, S. A. DeVita, Hellman, and Rosenberg’s cancer : principles & practice of oncology; Wolters Kluwer, 2019.

48. Vail, D. M. Withrow and MacEwen’s small animal clinical oncology; Elsevier, 2020.

(49) Holland, C. H.; Szalai, B.; Saez-Rodriguez, J. Transfer of regulatory knowledge from human to mouse for functional genomics analysis. Biochim Biophys Acta Gene Regul Mech 2020, 1863 (6), 194431. DOI: 10.1016/j.bbagrm.2019.194431 From NLM Medline.

(50) Garcia-Alonso, L.; Holland, C. H.; Ibrahim, M. M.; Turei, D.; Saez-Rodriguez, J. Benchmark and integration of resources for the estimation of human transcription factor activities. Genome Res 2019, 29 (8), 1363–1375. DOI: 10.1101/gr.240663.118.

(51) Reike, M. J.; de Jong, J. J.; Bismar, T. A.; Boorjian, S. A.; Mian, O. Y.; Wright, J. L.; Dall’Era, M. A.; Kaimakliotis, H. Z.; Lotan, Y.; Boormans, J. L.;, et al. Alignment of molecular subtypes across multiple bladder cancer subtyping classifiers. Urol Oncol 2024, 42 (6), 177 e175–177 e114. DOI: 10.1016/j.urolonc.2024.01.027.

(52) Alvarez, M. J.; Shen, Y.; Giorgi, F. M.; Lachmann, A.; Ding, B. B.; Ye, B. H.; Califano, A. Functional characterization of somatic mutations in cancer using network-based inference of protein activity. Nature Genetics 2016, 48 (8), 838–+. DOI: 10.1038/ng.3593.

(53) Badia, I. M. P.; Velez Santiago, J.; Braunger, J.; Geiss, C.; Dimitrov, D.; Muller-Dott, S.; Taus, P.; Dugourd, A.; Holland, C. H.; Ramirez Flores, R. O.;, et al. decoupleR: ensemble of computational methods to infer biological activities from omics data. Bioinform Adv 2022, 2 (1), vbac016. DOI: 10.1093/bioadv/vbac016.

(54) Bankhead, P.; Loughrey, M. B.; Fernandez, J. A.; Dombrowski, Y.; McArt, D. G.; Dunne, P. D.; McQuaid, S.; Gray, R. T.; Murray, L. J.; Coleman, H. G.;, et al. QuPath: Open source software for digital pathology image analysis. Sci Rep 2017, 7 (1), 16878. DOI: 10.1038/s41598-017-17204-5 From NLM Medline.

