## Supplementary Information for "Convergent biology, divergent drivers: a cross-species comparison of human and canine invasive urothelial carcinoma"

### SUPPORTING INFORMATION

**Supplementary Note 1:** Variance Decomposition Framework for Cross-Platform Comparisons

**S1.1 Sources of transcriptional variance in MIBC.** Comparing transcriptomes across species or platforms is an accounting problem: gene expression varies for several distinct reasons, and a fair comparison requires knowing which one a given difference reflects. The steady-state abundance of a transcript is set by a few largely separable inputs, written informally as

$$\Delta mRNA = f(DNA, epigentics, environment) + noise$$

The epigenetic term organizes bladder biology. The urothelium is a stratified epithelium with a fixed differentiation axis: basal cells rest on the basement membrane and remain proliferative and stem-like, while cells moving toward the lumen differentiate into superficial umbrella cells that express uroplakins and carry out the barrier and secretory functions of the luminal surface. Luminal and basal are first of all positions along this gradient, an epithelial developmental program conserved across mammals, which is why the same axis is recognizable in dog and human. MIBC inherits it as two molecular subtypes, luminal and basal, the major non-species source of transcriptional variance in these cohorts.

Cancer adds a term normal tissue lacks: somatic mutation. The two subtypes carry characteristic drivers upstream of the programs that enforce each state (Fig. S1A,B). FGFR3 marks the luminal subtype in human MIBC; BRAF V595E is the analogous luminal-associated driver in canine MIBC; NRF2, TP53, and RB1 mark the basal subtype. Each driver feeds a transcription-factor network that reinforces its differentiation state, GATA3-FOXA1-KLF4 in luminal tumors and STAT3-GATA6-HIF1α in basal tumors (Fig. S1A), with basal the more stem-like and EMT-primed of the two. The heatmap annotations (Fig. S1B) show this driver-subtype association holds in both species even though the drivers differ as genes.

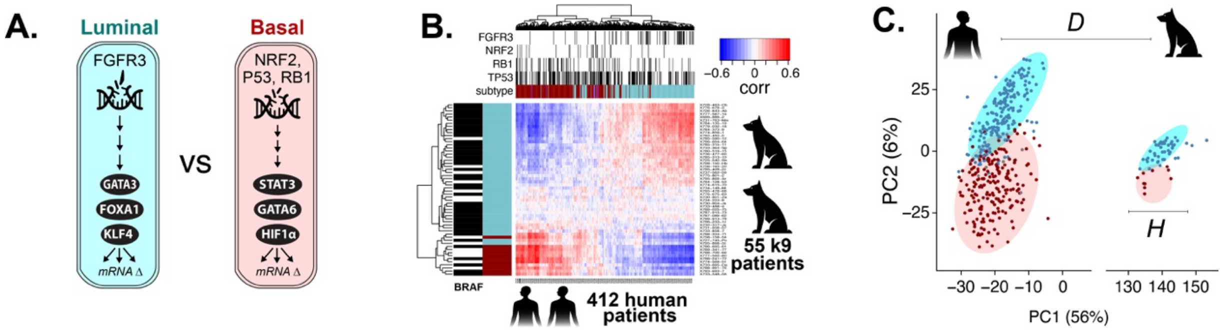


**Figure S1. Summary of mRNA and DNA variation in MIBC across human and canine patients.** A. conceptual summary of mutations and downstream transcription factors B. Corelation matrix visualizing molecular subtypes based on mRNA vectors and genetic variants C. PCA plot visualizing major sources of variance between datasets

In a combined cohort, then, the DNA term carries a large between-species component, the epigenetic term is dominated by the luminal-basal axis, the environment term becomes central later when tumors are compared to organoids, and noise is always present. The next section reorganizes these contributions into a decomposition that can be read off the data.

**S1.2 Variance decomposition for Cross-species comparison**. PCA of human (n = 408) and canine (n = 56) MIBC transcriptomes resolves two dominant axes (Fig. S1C): PC1 separates the species, PC2 separates luminal from basal within each species. We read this geometry as a decomposition of total observed variance,

$$\mathrm{Var}_{\mathrm{total}} = S + D + H + N$$

where S is signal conserved across species, D is between-species (and platform) divergence, H is within-species inter-patient heterogeneity, and N is technical noise. This is a conceptual scaffold, not a formal model with orthogonality assumptions. Each term reads off Fig. S1C: centroid separation along PC1 is

D, within-cluster spread is H + N, and the parallel orientation of the two clouds is S, the luminal-basal organization preserved in both species. S and D are the conserved and divergent halves of the DNA-and-development account, split at the species boundary; H is the within-species spread, dominated by the luminal-basal axis.

Both D and H mix biological and technical contributors that must be acknowledged before they can be separated. D can be genuine pathway-wiring divergence or a platform artifact such as library-preparation and normalization differences between cohorts. H can be real patient-tumor variation such as the luminal-basal axis, or technical sampling differences such as biopsy site and tumor purity. Disentangling them is the work of the sections that follow.

**S1.3 Correlation as a ratio of variance components.** The same decomposition makes the meaning of cross-species correlation explicit. When transcriptomic similarity between species is quantified by Pearson correlation, the squared correlation coefficient reflects the fraction of total observable variance attributable to conserved biology:

$$\boldsymbol{R^2 = \frac{S}{S + D + H + N}}$$

$R^2$ is therefore not a free-standing measure of species similarity. It is a ratio whose numerator is the biology of interest and whose denominator is inflated by every source of unwanted variation, biological or technical. A low cross-species $R^2$ can reflect genuine pathway divergence, or it can reflect platform artifacts; a high $R^2$ requires that both species' measurements be dominated by the same conserved program. The pairwise correlation structure in Fig. S1B should be read in this light: each cell of the heatmap is one realization of this ratio, and the block structure that emerges across patients and species reflects the relative magnitudes of S, D, and H in that comparison.

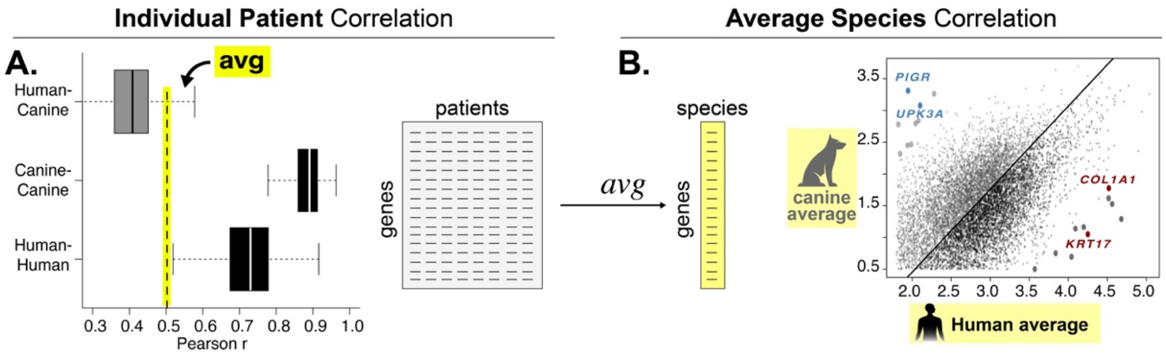


**Figure S2.** Comparison of individual patient correlation across species and average-patient correlation across species

Examination of the distribution of cross-species patient correlations (Fig. S2A) shows that while the conserved program S is present in the data, it never dominates any single cross-species pair. Individual human-canine correlations are low, because in each pair the denominator carries the full species offset (D) together with both tumors' independent heterogeneity (H) and noise (N). Within-species correlations are far higher, since same-species pairs share a common baseline and often the same subtype: the canine distribution sits highest, consistent with that cohort's uniform, luminal-dominated composition, and the human distribution lower, reflecting its greater subtype heterogeneity. These within-species values set the ceiling against which any cross-species comparison is measured.

**S1.4 Averaging to Reduce Noise and Patient Heterogeneity.** The variance decomposition implies a strategy for improving cross-species inference: reduce the denominator of $R^2$ by suppressing the variance components that obscure conserved biology. Averaging across n biological samples, whether across patients in bulk data or across cells in pseudo-bulked single-cell data, suppresses within-species heterogeneity and technical noise by a factor proportional to n, while the systematic terms S and D are unaffected:

$$R^2_{\text{avg}} = \frac{S}{S + D + H/n + N/n}$$

As n grows, H/n and N/n vanish and the cross-species $R^2_{avg}$ approaches S / (S + D), its ceiling now set by divergence alone. This is the species averaging of Fig. S1E,F: human and canine averages correlate strongly, most genes falling along the diagonal of concordant expression (S made visible), and the cross-species value rises from the individual-pair cloud toward the within-species range (the avg marker in Fig. S2A).

The residual divergence after averaging is itself informative. The genes that deviate most from the diagonal in Fig. S1F are not random species differences: canine-skewed outliers (PIGR, UPK3A) are luminal markers and human-skewed outliers (COL1A1, KRT17) are basal and stromal markers. Their apparent divergence reflects the differing prevalence of luminal and basal tumors in the two cohorts, and thus likely reflects sub-type bias (more luminal in canine) in patient-heterogeneity frequency.

**S1.5 Costs and Benefits of Averaging.** Aggregation is not free. Averaging across n samples reduces noise but also collapses biologically meaningful heterogeneity. This is desirable when *H* is dominated by sampling artifacts such as biopsy site or tumor purity, but destructive when *H* contains the inter-patient variation that precision medicine is trying to capture, such as the luminal-basal axis itself. Pathway aggregation has a parallel trade-off (next section): averaging across p genes within a well-curated pathway recovers coordinated program activity with high fidelity, but aggregation over poorly defined or heterogeneous gene sets averages real biology together with noise and can erase the signal it was meant to recover. The appropriate level of aggregation therefore depends on the question being asked, and the paired comparisons in the remainder of this work are designed to test cross-species fidelity at multiple levels: species-averaged programs, patient-resolved subtypes, and cell-type-resolved compositions.

**Supplementary Note 2:** Gene Signatures: suppress noise + while maintaining patient heterogeneity

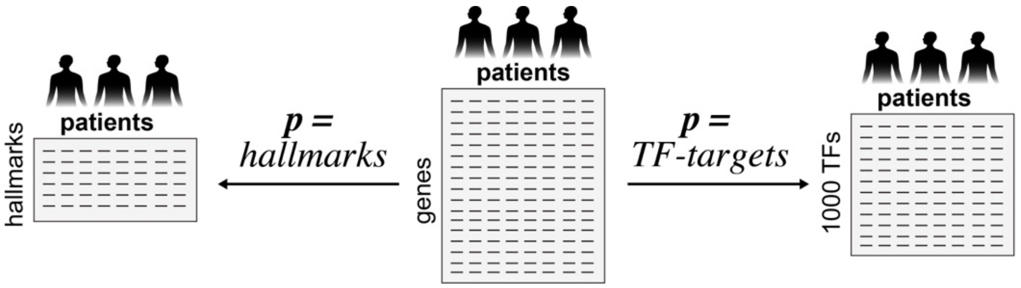


**Figure S3: Two major classes of "pathway" analysis applied to RNAseq datasets:** #1 statistically defined gene-sets defined by co-correlated transcriptional programs (e.g. MSigDB's Hallmark signatures), #2 Molecular Mechanism defined genesets defined by the bone fide transcriptional targets of transcription factors (e.g. ChIPseq, motif binding, etc.)

Gene signatures replace the high-dimensional gene-by-patient matrix with a lower-dimensional score matrix by aggregating the expression of the *p* genes that mark a common process or cell type, computed independently within each patient (Fig. S3, top). The same operation underlies all three signature methods used here: a pathway or hallmark score pools the genes of a transcriptional program, a transcription-factor activity score pools a regulator's downstream targets, and a cell-type deconvolution estimate pools a cell type's marker genes. In each case one patient's high-noise readout across many genes is collapsed into a single, more stable number.

In the decomposition of Note S1, this aggregation acts on the noise term alone. The *p* genes of a signature carry largely independent technical noise but share the conserved program *S* and respond coherently to a given patient's state, so pooling them suppresses *N* by roughly a factor of *p* while leaving *S*, *D*, and the inter-patient heterogeneity *H* intact:

$$R^2_{\text{sigs}} = \frac{S}{S + D + H + N/p}$$

This is the complement of the patient averaging in Note S1, $R^2_{avg} = S / (S + D + H/n + N/n)$, which suppressed *H* together with *N*. Patient averaging gains cross-species signal by discarding inter-patient differences; signature scoring gains it by cleaning up each patient's measurement while keeping those differences. The *H* term is retained deliberately, because inter-patient heterogeneity, the luminal-basal axis and the composition of the microenvironment, is the biology these analyses are meant to compare, not a nuisance to be averaged away.

Scoring human and canine tumors on shared signatures shows the effect directly (Fig. S3). Cross-species correlation is high for both transcription-factor activity (r = 0.917) and pathway hallmarks (r = 0.932), with conserved programs such as proliferation (MYC, G2M, mitotic spindle), protein secretion, and oxidative phosphorylation falling on the diagonal and only a few interpretable signatures off it. In the correlation-context panels, the cross-species ("cross-condition") distribution sits just below the within-species inter-sample distributions rather than far beneath them: the conserved signal now approaches the reproducibility ceiling set by the residual *H* and *D*. This is the same *S* that was buried in the gene-level cross-species correlations of Fig. S1D, brought to the surface by collapsing *N* rather than by averaging patients together (Fig S3).

Because *H* survives, signature scores still resolve individual patients, separating luminal from basal tumors and quantifying microenvironmental composition within each species, while reporting a low-noise, cross-species-comparable readout of known biology. Signature scoring and patient averaging are therefore two aggregations chosen for different ends: averaging over patients (*n*) to ask whether the two species share a program at all (Fig. S1), and averaging over genes within a signature (*p*) to compare that program patient-by-patient across species, the basis for the pathway and deconvolution results of Fig. 2.

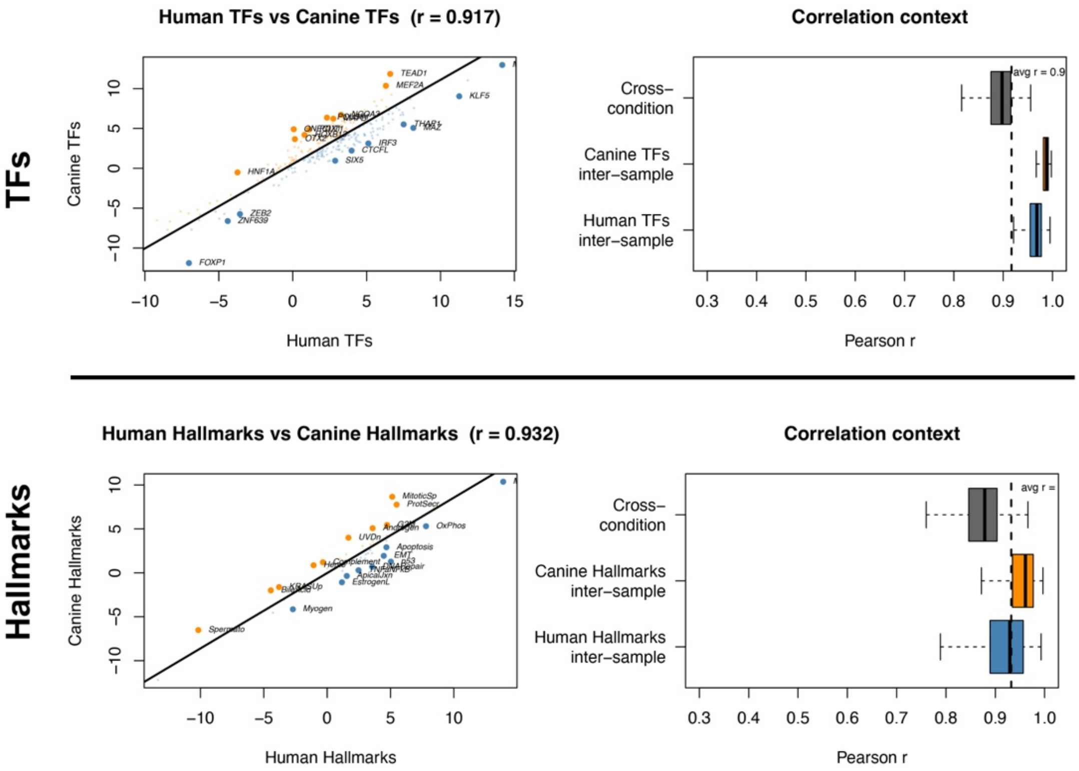


Figure S4. Comparison of average and individual patient correlations in "pathway transformed" mRNA space
